# *In vitro* and *in silico* modeling of cellular and matrix-related changes during the early phase of osteoarthritis

**DOI:** 10.1101/725317

**Authors:** Marie-Christin Weber, Lisa Fischer, Alexandra Damerau, Igor Ponomarev, Moritz Pfeiffenberger, Timo Gaber, Sebastian Götschel, Jens Lang, Susanna Röblitz, Frank Buttgereit, Rainald Ehrig, Annemarie Lang

## Abstract

**Objective:** Understanding the pathophysiological processes of osteoarthritis (OA) require adequate model systems. Although different *in vitro* or *in vivo* models have been described, further comprehensive approaches are needed to study specific parts of the disease. This study aimed to combine *in vitro* and *in silico* modeling to describe cellular and matrix-related changes during the early phase of OA. We developed an *in vitro* OA model based on scaffold-free cartilage-like constructs (SFCCs), which was mathematically modeled using a partial differential equation (PDE) system to resemble the processes during the onset of OA.

**Design:** SFCCs were produced from mesenchymal stromal cells and analyzed weekly by histology and qPCR to characterize the cellular and matrix-related composition. To simulate the early phase of OA, SFCCs were treated with interleukin-1β (IL-1β), tumor necrosis factor α (TNFα) and examined after 3 weeks or cultivated another 3 weeks without inflammatory cytokines to validate the regeneration potential. Mathematical modeling was performed in parallel to the *in vitro* experiments.

**Results:** SFCCs expressed cartilage-specific markers, and after stimulation an increased expression of inflammatory markers, matrix degrading enzymes, a loss of collagen II (Col-2) and a reduced cell density was observed which could be partially reversed by retraction of stimulation. Based on the PDEs, the distribution processes within the SFCCs, including those of IL-1β, Col-2 degradation and cell number reduction was simulated.

**Conclusions:** By combining *in vitro* and *in silico* methods, we aimed to develop a valid, efficient alternative approach to examine and predict disease progression and new therapeutic strategies.

## Introduction

The pathogenesis of osteoarthritis (OA) has not been fully understood so far, partly due to the lack of an optimal model system which sufficiently incorporates all aspects of the disease [1, 2]. A vast variety of *in vivo, ex vivo, in vitro* and (to some extent) *in silico* models for OA already exists, and these all try to mimic the main features of OA pathophysiology including the degradation of extracellular matrix (ECM), inflammation and alterations in cell metabolism, viability and differentiation [3-5].

*In vivo* OA models are crucial for translational research, but mainly make use of small rodents, especially mice, although these species show significant differences in articular cartilage anatomy, loading conditions and life span [6, 7]. Thus, *in vitro* models represent an important tool to investigate the mechanisms involved in the pathogenesis of OA and to examine possible therapeutic values. Model systems include monolayer cultures with cell lines or primary chondrocytes, co-cultures, 3D-cultures and cartilage explants from either humans or animals [8]. However, a major challenge is the source of primary chondrocytes or explants, since healthy human cartilage samples are rare, and thus sample collection is mainly undertaken from total joint replacement surgeries. Independent of the cell source, it is a consensus that the 3D cultivation of primary chondrocytes resembles the *in vivo* situation more closely. Most 3D culture systems involve a scaffold to provide the cells with a predetermined structure where their distinct effects on the cells are often not considered. Recent developments aim to establish scaffold-free tissue engineered cartilage mainly from human chondrocytes which could serve as a promising approach to cartilage repair *in vivo*, as well as an excellent *in vitro* model, since ECM formation and degradation can be evaluated without the interference of a scaffold [9-11].

In addition to biological models, *in silico* modeling offers a powerful tool to bring clarity to the processes that evolve during *in vitro* or *in vivo* modeling [12, 13]. Few studies have been performed so far to obtain parameters for *in vitro* modeling, although a combination of biological data with mathematical modeling promises to accelerate translation in OA research [14-16]. Catt et al. describe a partial differential equation (PDE) model of cartilage with a focus on the production of ECM and the growth of chondrocytes synergistically leading to tissue expansion [15]. Moreover, Kar et al. investigate cartilage degradation induced by interleukin-1β (IL-1β) using a PDE model based on data from the literature together with existing experimental data to calibrate their model [16]. In contrast, Baker et al. use an ordinary differential equation (ODE) system to describe the interaction of anti-inflammatory cytokines, proteinases and fibronectin in arthritic cartilage [17]. Since biomechanical influences are of utmost interest, several approaches here have already been used, and these include model arthritic cartilage under cyclic compressive loading or a combining of inflammation and biomechanics [18, 19].

However, most of the above-mentioned models and approaches have been developed almost independently from experiments lacking the combination of *in vitro* and *in silico* modeling processes. Thus, the objective of our study was to develop a human *in vitro* OA model based on engineered tissue and scaffold-free cartilage-like constructs (SFCCs) sourcing from human mesenchymal stromal cells (hMSCs). These SFCCs were mathematically modeled in parallel based on PDEs resembling the matrix degradation processes during the early phase of OA.

## Material & Methods

### General study design

**Figure 1:**
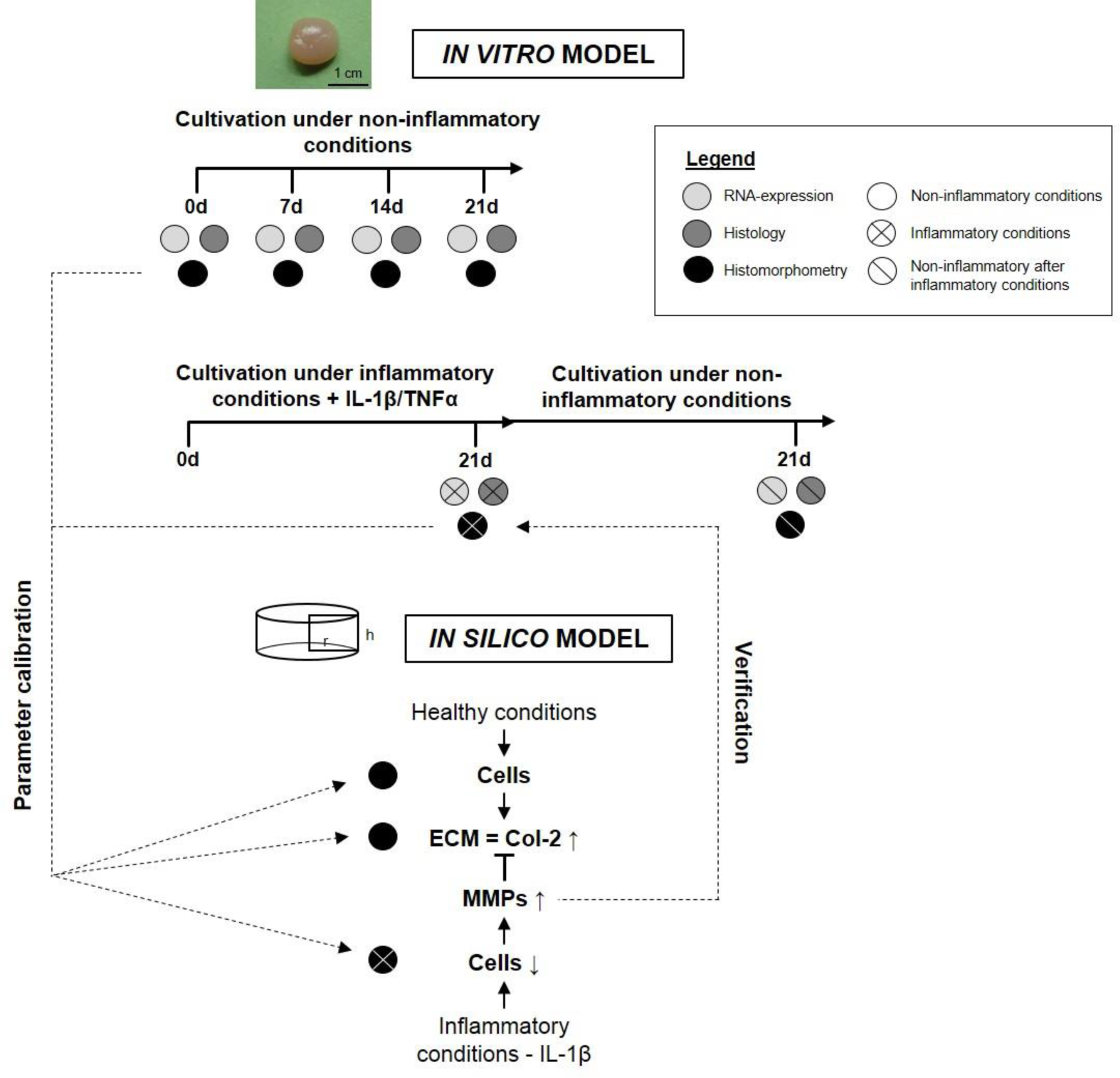
Overview on our current approach and study design.

### hMSC isolation and expansion

Human mesenchymal stromal cells (hMSCs) were obtained from human femoral bone marrow of patients undergoing hip arthroplasty (provided by the Centre for Musculoskeletal Surgery, Charité – Universitätsmedizin Berlin and distributed by the “Tissue Harvesting” Core Facility of the Berlin Brandenburg Centre for Regenerative Therapies, Berlin, Germany). All protocols were approved by the Charité – Universitätsmedizin Ethics Committee and performed according to the Helsinki Declaration (ethical approval EA1/012/13). Donor information is summarized in Table 1. hMSCs were cultivated in DMEM GlutaMAX™ Medium (Thermo Fisher Scientific, MA) with 20 % (v/v) StemMACS™ (Miltenyi Biotech, Germany), 10 % (v/v) fetal calf serum (FCS) (Thermo Fisher Scientific, MA), 100 units/ml penicillin and 0.1 mg/ml streptomycin (Thermo Fisher Scientific, MA) at a temperature of 37 °C in 5 % CO_2_ atmosphere. hMSCs were cultured separately for each donor and characterized by flow cytometry (CD90^+^, CD105^+^, CD73^+^, CD14^−^, CD20^−^, CD34^−^, CD45^−^, HLA-DR^−^) as well as osteogenic, adipogenic and chondrogenic differentiation assays. Only cells successfully passing the characterization were further cultivated up to passages 3-5.

**Table 1:** hMSC donor information and related experiments.

| Patient | Age | Sex | Characterization | Type of experiment |
| --- | --- | --- | --- | --- |
| 1 | 71 | female | + | Experiments under<br>non-inflammatory conditions<br>Histology<br>RNA analysis |
| 2 | 56 | female | + |  |
| 3 | 85 | female | + |  |
| 4 | 59 | female | + | Experiments under<br>inflammatory conditions<br>Histology<br>RNA analysis |
| 5 | 79 | male | + |  |
| 6 | 66 | male | + |  |

### Generation of SFCCs and experimental setup

SFCCs were produced based on a patented protocol [20]. In short, around 10-15 million hMSCs were transferred into a 3D state via centrifugation and self-assembly followed by maturation for 3 to 4 weeks applying biomechanical forces until reaching cartilage-like stiffness [21]. In advance of the experiments, SFCCs were cultured for 4 weeks in DMEM GlutaMAX™ Medium, supplemented with 10 % (v/v) FCS, 100 units/ml penicillin and 0.1 mg/ml streptomycin (all from Thermo Fisher Scientific, MA) and 9.39 mg/l ascorbic acid (Sigma-Aldrich, MO); in the following, this is referred to as ‘standard medium’. In a first step, SFCCs were cultivated under cytokine-free conditions and samples were taken on days 0, 7, 14 and 21 (n = 3 per time point). Secondly, to resemble the proinflammatory state that has been described in patients with early OA, stimulation was performed using standard medium supplemented with 50 ng/ml recombinant human IL-1β (2 × 10^8^ IU/mg) and 100 ng/ml recombinant human tumor necrosis factor alpha (TNFα) (2 × 10^7^ IU/mg; both from ImmunoTools, Germany). Samples of cytokine stimulated SFCCs (STIM) and controls (CTL) were taken on day 21 (n = 9 with triplicates of 3 different donors). A subset of cytokine stimulated SFCCs was further cultivated for 21 more days under cytokine free conditions to evaluate whether regeneration might occur (REG) (n = 9 with triplicates of 3 different donors). SFCCs from each donor were randomized into groups and analysis was performed in a blinded fashion by numbering samples at random.

### Histological staining and immunofluorescence

Samples for histological analysis were first fixed in 4 % paraformaldehyde (PFA) for 6 h and subsequently treated with 10%, 20% and then 30% glucose solution, each for 24 h. After fixation, samples were cryo-embedded in SCEM embedding medium and cryo-sections of 8-µm thickness were prepared using cryofilms (Sectionlab, Japan). Prior to each histological and immunohistochemical staining, procedure slices were dried for 20 min at room temperature.

### H&E and Alcian blue staining

Hematoxylin and eosin (H&E) staining was performed according to the following protocol: fixation with 4% PFA (10 min), washing with distilled water (5 min), first staining step in Harris’s hematoxylin solution (7 min) (Merck, Germany), washing with distilled water (2x), differentiation step in 0.25 ml of concentrated HCl in 100 ml of 70 % ethanol, washing with tap water (2× 10 min), second staining step in 0.2% eosin (2 min) (Chroma Waldeck, Germany), differentiation in 96% ethanol, washing in 96% ethanol, 100% ethanol (2× 2 min), fixation with xylol (2× 2 min), covering of stained slices with Vitro-Clud^®^ (R. Langenbrinck GmbH, Germany). Alcian blue staining was performed according to the following protocol: fixation with 4% PFA (10 min), washing with distilled water (5 min), 3% acetic acid (3 min), first staining step in 1% Alcian blue 8GX (Sigma-Aldrich, MO) in 3% acetic acid, pH 2.5 (30 min), washing in 3% acetic acid, washing in distilled water, second staining step in Nuclear fast red-aluminum sulfate solution (Chroma Waldeck, Germany), washing in distilled water, graded ethanol series (80%, 96%, 100%) (2 min each), fixation with xylol (2x 2 min), covering of stained slices with Vitro-Clud^®^ (R. Langenbrinck GmbH, Germany).

### Immunohistochemistry

Immunohistochemistry was performed according to the following protocol: rehydration with phosphate buffered saline (PBS) (10 min), blocking with 3% H_2_O_2_ (30 min), washing with PBS (5 min), blocking with 5% normal horse serum (Vector Laboratories, CA) in 2% bovine serum albumin (BSA) / PBS, overnight incubation with primary antibody for collagen type I and collagen type II at 4 °C (Ms mAb to collagen I, ab6308, 1:500, Abcam, UK) (collagen type II Ms 6B3, 1:10, quartett Immunodiagnostika, Germany), washing in PBS (2x 5 min), incubation with 2 % secondary antibody (biotinylated horse anti-mouse IgG antibody, Vector Laboratories, CA) diluted in 2x normal horse serum / 2% BSA / PBS (30 min), washing in PBS (2x 5 min), incubation with avadin-biotin complex (VECTASTAIN^®^ Elite^®^ ABC HRP Kit, Vector Laboratories, CA) (50 min), washing with PBS (2× 5 min), incubation with DAB under microscopic control with time measurement (DAB peroxidase (HRP) Substrate Kit, Vector Laboratories, CA), stopping with PBS (2x), washing in distilled water, counterstaining in Mayer’s hematoxylin 1:2 (Sigma-Aldrich, MO), blueing in tap water (5 min), washing in distilled water, covering of stained slices with Aquatex^®^ (Merck, Germany). Pictures were taken with the Axioskop 40 optical microscope (Zeiss, Germany) with AxioVision microscopy software (Zeiss, Germany).

### Immunofluorescence staining – MMP1 and MMP13

Immunofluorescence staining was used to quantify the MMPs. First of all, the slides were air-dried at room temperature and then rehydrated with PBS for 10 min. Subsequently, unspecific binding sites were blocked with PBS/5% FCS for 30 min. Primary MMP13 antibody (mouse anti-human; Invitrogen Thermo Fisher Scientific, monoclonal, MA5-14247) was diluted 1:200 in PBS/5% FCS/0.1% Tween^®^ 20 and primary MMP1 antibody (mouse anti-human; Invitrogen Thermo Fisher Scientific, monoclonal, MA5-15872) was diluted 1:500 in PBS/5% FCS/0.1% Tween^®^ 20 and incubated according to the manufacturer’s instruction for 3 h. After each incubation step, the preparation was washed 3 times with PBS/0.1% Tween^®^ 20. Secondary antibody (goat anti-mouse A546; Invitrogen Thermo Fisher Scientific, A-11003) was diluted 1:500 in PBS/5% FCS/0.1% Tween^®^ 20 and applied for 2 h. In the final staining step, core staining was performed using DAPI (1 µg/ml diluted in PBS/5% FCS/0.1% Tween^®^ 20) for 15 min. After air bubble-free covering with FluoroMount covering medium, microscopic evaluation was performed with the fluorescence microscope BZ-9000A (Keyence, Germany) using the DAPI and TRITC channels. Image analysis was performed using ImageJ. Determining the cell number the find maxima tool was used for the DAPI image, whereas the area of MMP positive signals was measured using the color threshold tool.

### Histomorphometry

Histomorphometry was performed using FIJI ImageJ 1.52i [22, 23]. H&E stained sections were used to analyze the cell count per area and cell distribution within the SFCCs. The cell count per tissue area (cells/mm^2^) was identified from HE overview pictures of each SFCC with 50x magnification using a modified color deconvolution method [24] (for additional detail information see Fig. 2 and Table 2). First, a free hand selection tool was used to define the region of interest (ROI) for the section outline representing the total area (Tt.A.) of the section. The gap area (Gp.A.) where no tissue was present was identified using the Color Threshold tool and subtracted from the Tt.A. to obtain the Total Tissue Area (Tt.T.A.). Next, the Color Deconvolution plugin of ImageJ with a vector to separate hematoxylin and eosin staining into each color layer was applied. Within the hematoxylin layer, the cell nuclei were identified by applying the Threshold Tool of ImageJ based on their brightness within the layer. A binary image was created representing the nuclei of the cells within the section. Finally, cell count was performed from these binary images with a combination of the Particle Analysis tool in ImageJ and manual counting.

**Table 2:**
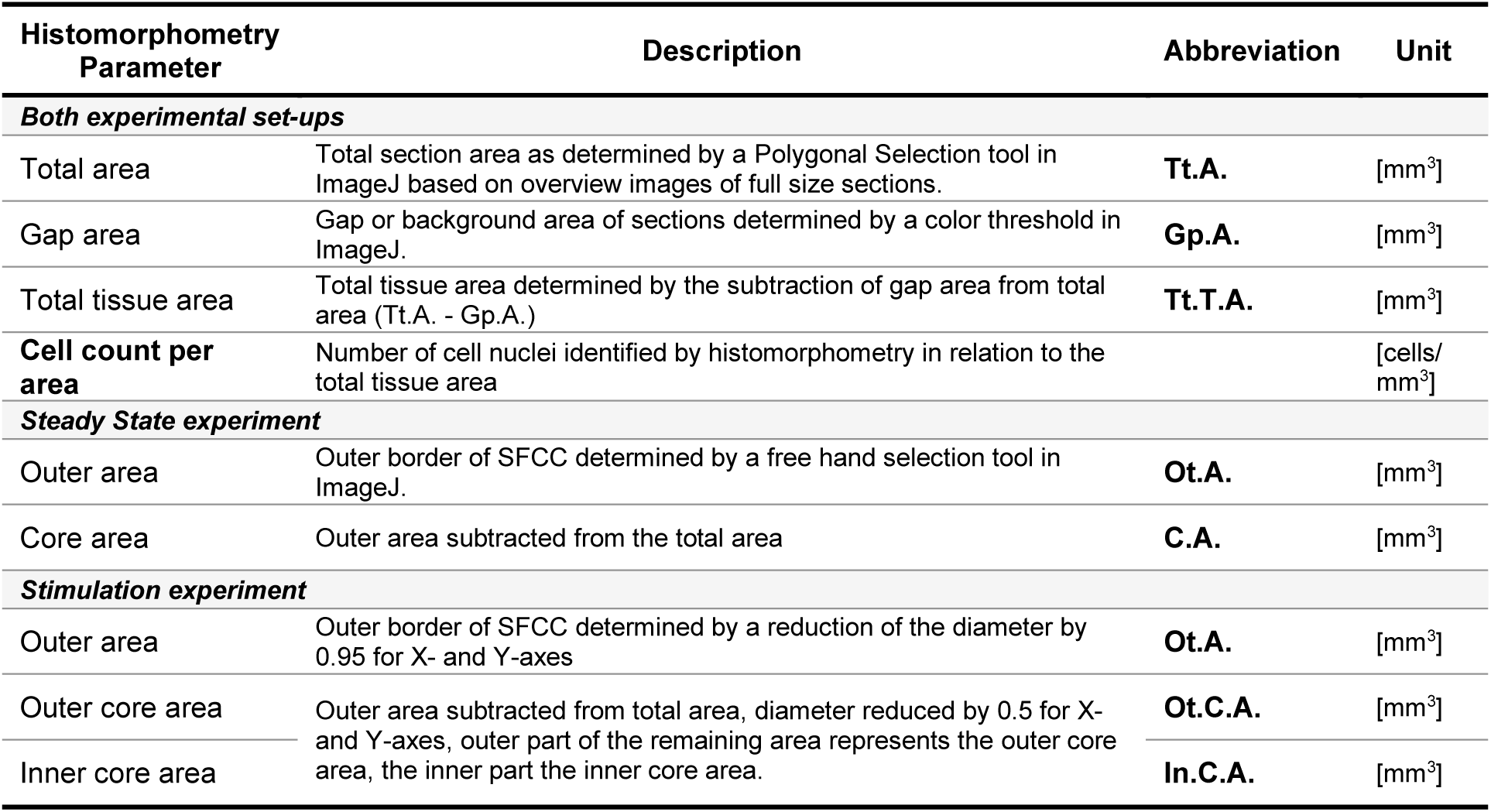
Histomorphometry parameters.

| Histomorphometry Parameter | Description | Abbreviation | Unit |
| --- | --- | --- | --- |
| **Both experimental set-ups** |  |  |  |
| Total area | Total section area as determined by a Polygonal Selection tool in ImageJ based on overview images of full size sections. | **Tt.A.** | [mm<sup>3</sup>] |
| Gap area | Gap or background area of sections determined by a color threshold in ImageJ. | **Gp.A.** | [mm<sup>3</sup>] |
| Total tissue area | Total tissue area determined by the subtraction of gap area from total area (Tt.A. - Gp.A.) | **Tt.T.A.** | [mm<sup>3</sup>] |
| **Cell count per area** | Number of cell nuclei identified by histomorphometry in relation to the total tissue area |  | [cells/mm<sup>3</sup>] |
| **Steady State experiment** |  |  |  |
| Outer area | Outer border of SFCC determined by a free hand selection tool in ImageJ. | **Ot.A.** | [mm<sup>3</sup>] |
| Core area | Outer area subtracted from the total area | **C.A.** | [mm<sup>3</sup>] |
| **Stimulation experiment** |  |  |  |
| Outer area | Outer border of SFCC determined by a reduction of the diameter by 0.95 for X- and Y-axes | **Ot.A.** | [mm<sup>3</sup>] |
| Outer core area | Outer area subtracted from total area, diameter reduced by 0.5 for X- and Y-axes, outer part of the remaining area represents the outer core area, the inner part the inner core area. | **Ot.C.A.** | [mm<sup>3</sup>] |
| Inner core area |  | **In.C.A.** | [mm<sup>3</sup>] |

**Figure 2:**
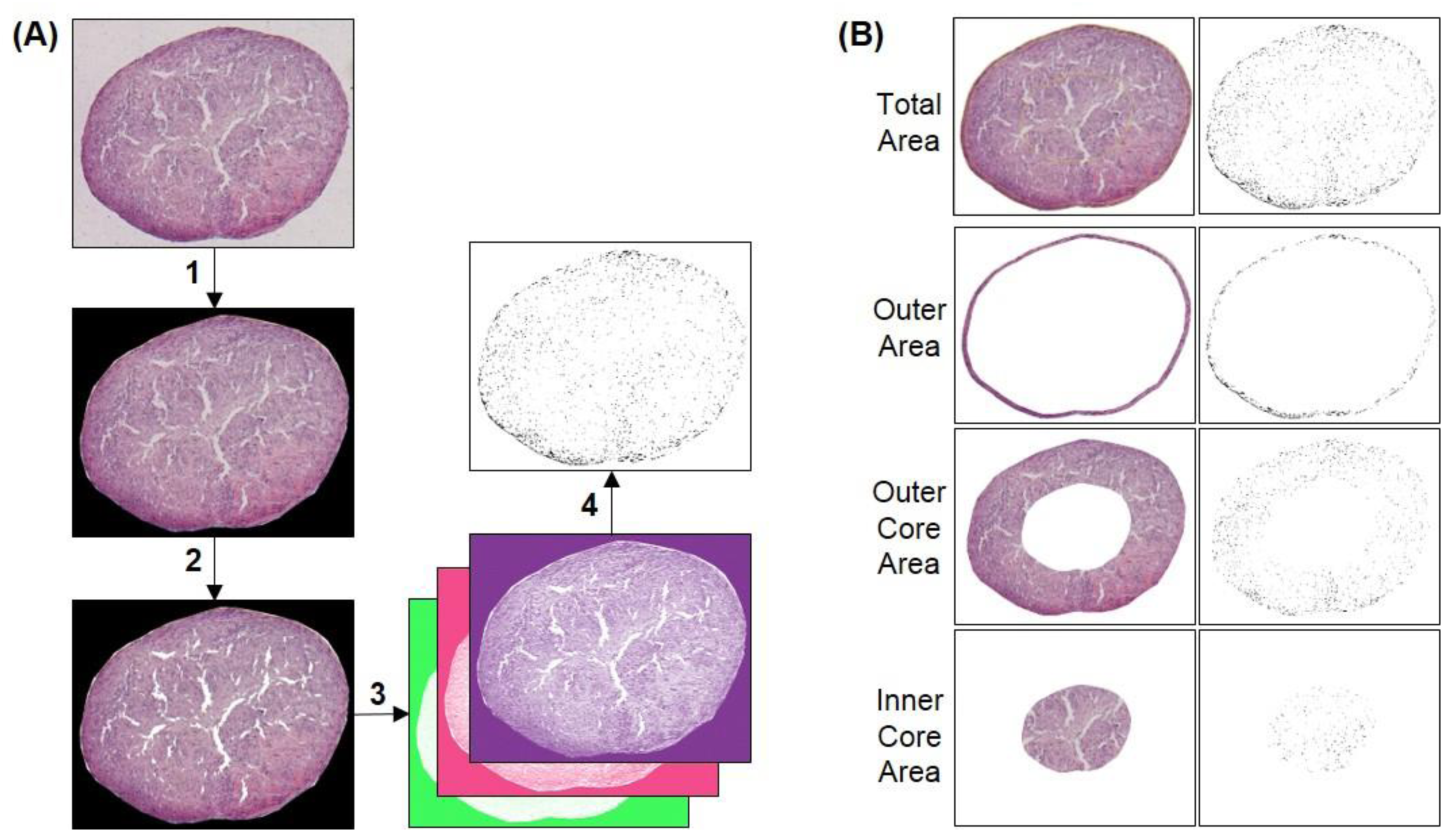
Pipeline for histomorphometric analysis. (**A**) Histomorphometric analysis for cell count per area using a modified color deconvolution method. A 50x magnification overview image of an H&E stained section was taken. The section outline was defined using the Polygonal Selection tool in ImageJ (1). In order to subtract the background (or gap area) the Color Threshold tool was utilized (2). Next, the Color Deconvolution plugin (3) was used to obtain the hematoxylin color channel only in which the cell nuclei could be identified using a threshold for brightness (4). Particle analysis could then be performed based on the binary image attained in the previous step to obtain the cell count per area [cells/mm2]. (**B**) For further analysis, the ‘Total Area’ was further divided into an ‘Outer Area’ by scaling the region of interest (ROI) for ‘Total Area’ to 0.95 for X- and Y-axes. The remaining ‘Core Area’ was then divided into an ‘Outer Core Area’ and an ‘Inner Core Area’ by scaling the ROI for the ‘Core Area’ by 0.5 for the X- and Y-axes.

**Figure 3:**
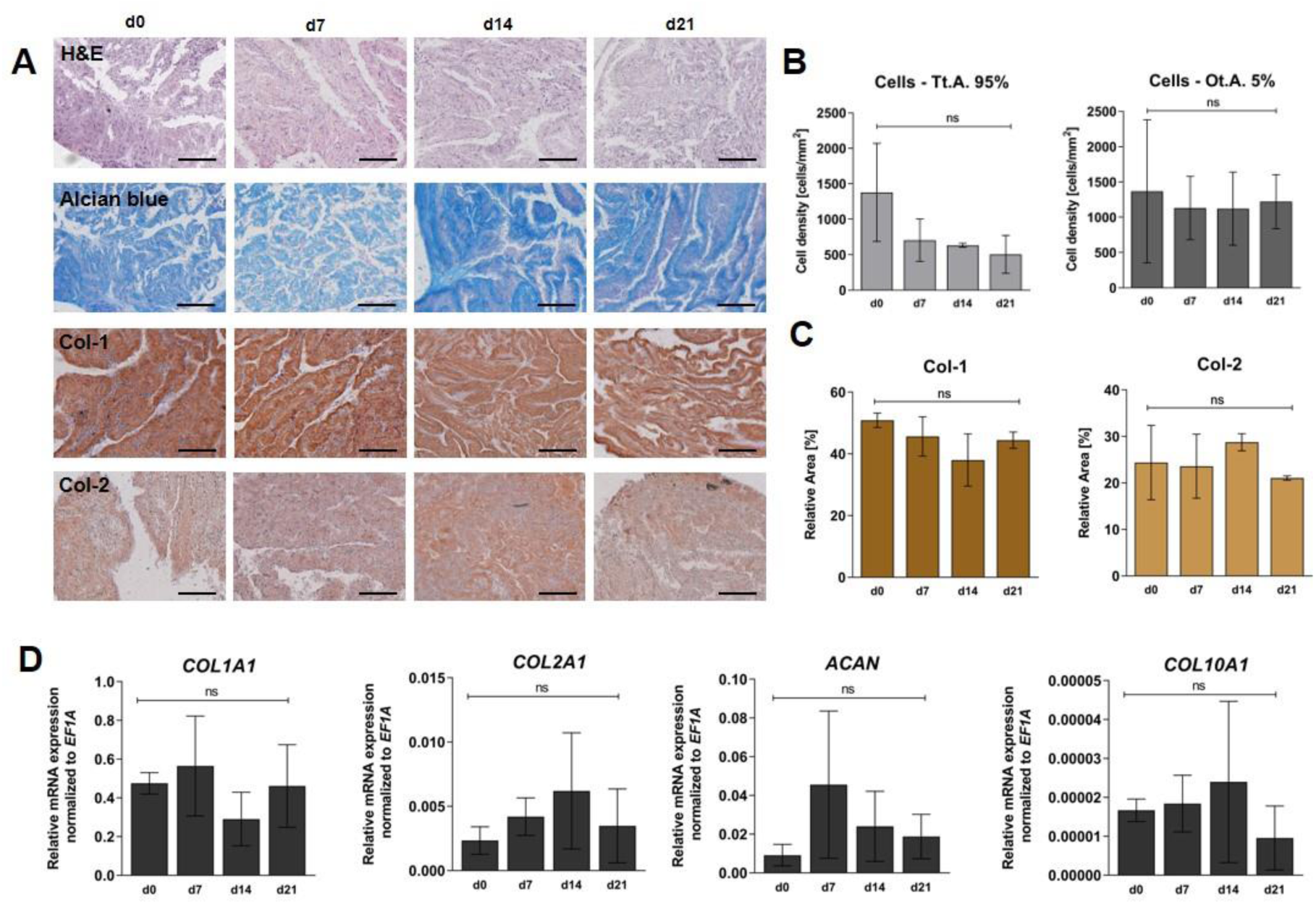
Experimental results after cultivating SFCC for 3 weeks under non-inflammatory conditions. (A) Exemplary images for histological staining with hematoxylin & eosin (H&E), Alcian blue, Col-1 and Col-2 immunohistochemistry, scale bars = 200 µm at 100x magnification. (B) Histomorphometry results for cell count within total area (Tt.A.) and outer area (Ot.A.). X-axes shows time points from d0-d21, y-axes show cell count [cells/m^3^], bars indicate mean ± SD. (C) Immunohistochemistry coverage for Col-1 and Col-2. X-axes shows time points from d0-d21, y-axes show relative coverage area in %, bars indicate mean ± SD. (D) Gene expression studied via qPCR for *COL1A1, COL2A1, COL10A1* and *ACAN*. X-axes shows time points from d0-d21, y-axes show relative mRNA expression normalized to the housekeeper *EF1A* graph bar with mean ± SEM. Statistical differences between groups was tested with the Kruskal-Wallis test and Dunn’s multiple comparisons test (see also Supplementary Results Table S1). ns = p > 0.05.

**Figure 4:**
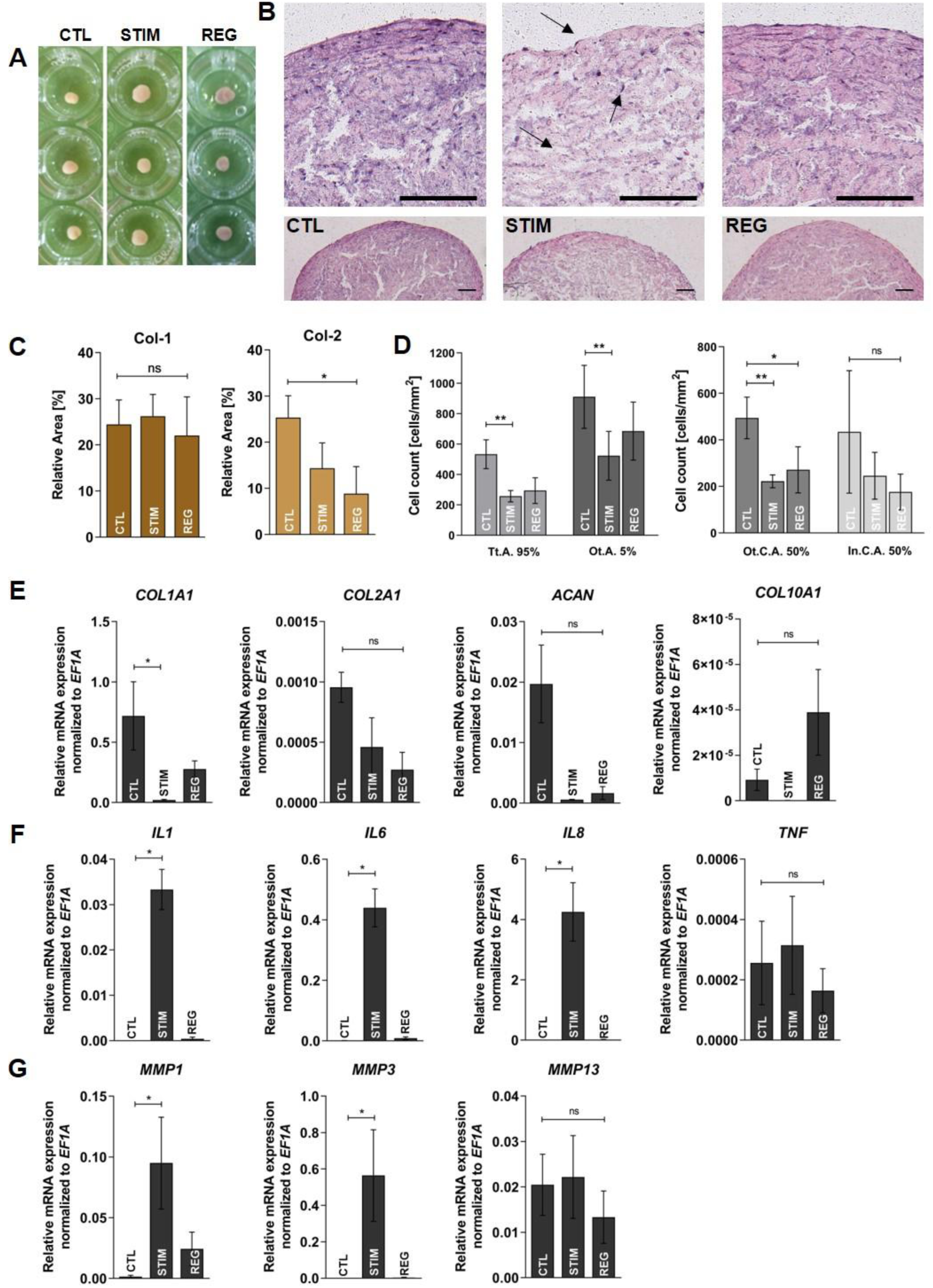
Experimental results after cultivating SFCC for 3 weeks under non-inflammatory conditions (CTL), with IL-1β and TNFα (STIM) or under non-inflammatory conditions for additional 3 weeks after stimulation (REG). (A) Macroscopic overview on different SFCCs exemplary for each condition. (B) Exemplary images of SFCCs stained with H&E for CTL, STIM and REG group. Upper row is 200x magnification, bottom row is 50x magnification, scale bars = 200 µm. Arrows indicate tissue softening and changes of cellular phenotypes. (C) Immunohistochemistry coverage for Col-1 and Col-2. X-axes show experimental groups, Y-axes show relative coverage area in %, bars indicate mean ± SD. (D) Histomorphometry results for cell count within total area (Tt.A.), outer area (Ot.A.), outer core area (Ot.C.A.) and inner core area (In.C.A.). X-axes shows time points from d0-d21, Y-axes show cell count [cells/m^3^], bars indicate mean ± SD. (E-G) Gene expression studied via qPCR for *IL1, TNF, IL6, IL8, MMP1, MMP3, MMP13, COL1A1, COL2A1, ACAN*. X-axes shows experimental groups, Y-axes show 2^-ΔCt normalized to the housekeeper *EF1A* depicted as graph bar with mean ± SEM. Statistical differences between groups were tested with the Kruskal-Wallis test and Dunn’s multiple comparisons test (see also Supplementary Results Table S2 and S3). ns = p > 0.05, *p < 0.05, **p < 0.01.

**Figure 5:**
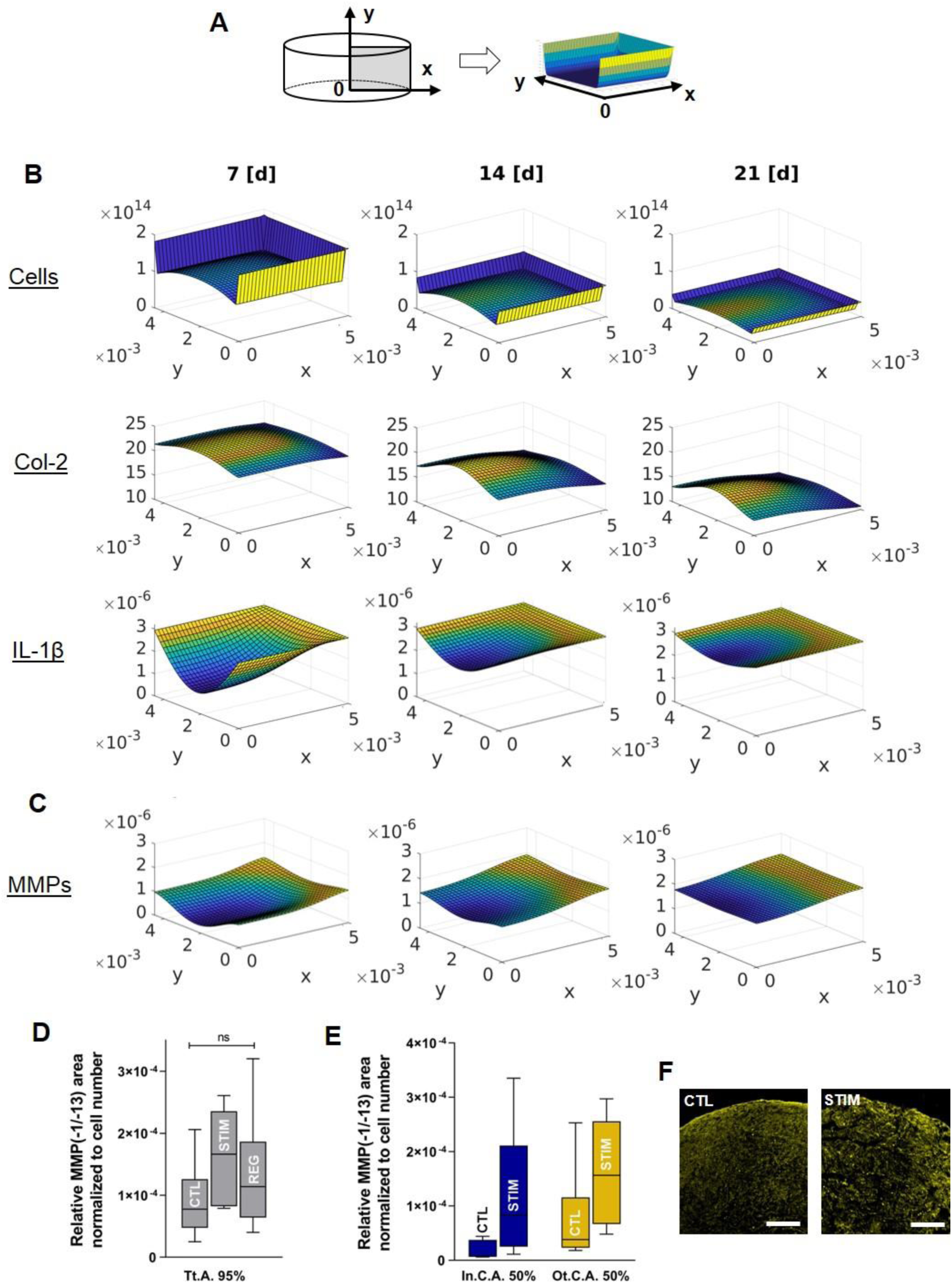
Results from the *in silico* model. (A) Explanation of the dimensions and axes. (B) Spatial-temporal development of Cell [1/m^3^] (first row), Col-2 [%] (second row), IL-1*β* [mol/m^3^] (last row) and (C) MMP [mol/m^3^] (third row) over 3 weeks. The PDE model described in Table 5 was solved with the adaptive finite element toolbox KARDOS [25, 26] and MATLAB. (D) MMP-1/-13 were stained in CTL, STIM and REG and normalized to the cell count (total area = Tt.Ar.). (E) Quantitative analysis was performed for inner core (In.C.A.) and outer core area (Ot.C.A.) for CTL and STIM are only to be compared to the results from the *in silico* model. Graphs show box and whiskers plots (Min-Max). (F) Exemplary images for MMP-1 staining CTL vs. STIM. The scale bar indicates 200 µm.

The cell count for the Tt.T.A. was performed identically for the experiments under normal and inflammatory conditions. A ROI for the Outer Area (Ot.A.) was identified with a manual selection tool for the experiment under normal conditions. For the stimulation experiment, ROIs were determined for the Ot.A. with a reduction of section diameter to 0.95 for X- and Y-axes, which then was further divided in an outer core area (Ot.C.A.) and inner core area (In.C.A.) by another reduction of the diameter by 0.5 for X- and Y-axes. Sections of 2 different levels per SFCC were analyzed respectively and the mean taken for statistical analysis. In order to analyze the immunohistochemically stained sections, the DAB area coverage was measured for Type 1 collagen (Col-1) and Type 2 collagen (Col-2), respectively. Two to three pictures of 100x magnification per section were analyzed and the mean taken for statistical analysis. Tt.T.A was determined again by measuring the Tt.A. and subtracting Gp.A. which was identified with the Color Threshold tool in ImageJ. The area stained positive for Col-1 and Col-2 was defined by the Color Threshold tool also. Color Thresholds were determined for each set of staining separately.

### RNA isolation, cDNA synthesis and qPCR

Total RNA was isolated from the SFCCs using TissueRuptor II (QIAGEN, Germany) to homogenize the tissue and the RNeasy Fibrous Tissue Mini Kit (QIAGEN, Germany) was used to extract the RNA according to the provided protocols. RNA concentrations were measured via NanoDrop Fluorometer (Thermo Fisher Scientific, MA) and RNA integrity was confirmed via the 2100 Bioanalyzer (Agilent Technologies, CA). Sensiscript RT Kit (QIAGEN, Germany) was used for cDNA synthesis with 50 ng per reaction according to the manufacturers’ instructions. Primers were designed using Primer Blast (NCBI, MD) and sequence analysis of qPCR products was performed at LGC genomics (LGS genomics GmbH, Berlin, Germany) to confirm primer specify (for primer sequences see Table 3). To analyze RNA expression, quantitative PCR (qPCR) was performed using the DyNAmo ColorFlash SYBR Green qPCR kit (Thermo Fisher Scientific, MA) at a Mx3000P qPCR System (Agilent Technologies, CA) with approximately 1.5 ng cDNA per 20 µl reaction and the following temperature profile: 7 min denaturation at 95 °C, 45 cycles of 5 × at 95 °C, 7 s at 57 °C and 9 s at 72 °C. Two technical replicas per sample and gene were performed. After each qPCR run, a melting curve analysis was performed to confirm primer specificity. In cases where no amplification curve reached the threshold before 45 cycles, the threshold-cycle value (Ct-value) was assumed to be 45. Gene expression data is shown as ΔCt-value (= 2^-ΔCt) with normalization to the housekeeper *elongation factor 1-alpha* (*EF1A*).

**Table 3:**
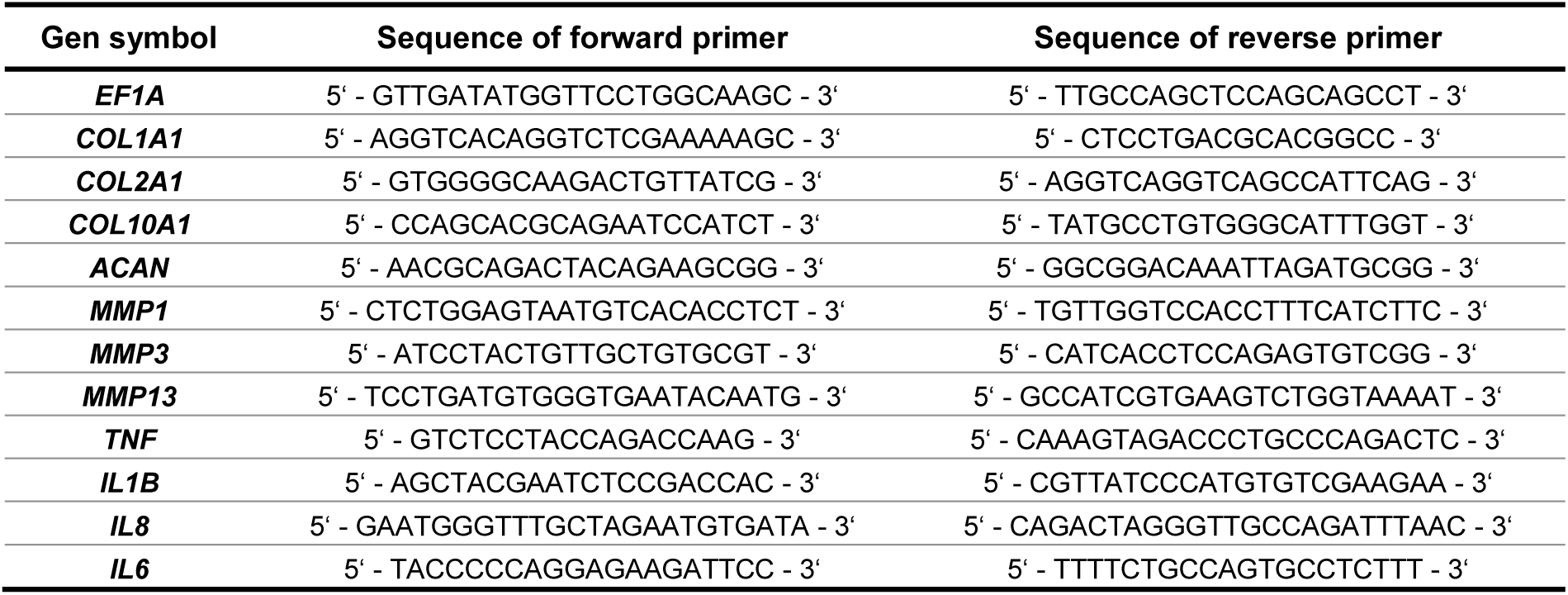
Sequences of primers used for qPCR.

### RNA isolation from human cartilage

Human cartilage was collected from femoral condyles taken during total knee replacement surgeries (ethical approval EA1/012/13). Cartilage was harvested from areas as unaffected as possible and transferred to RNAlater (QIAGEN, Germany) for 1 h at 4 °C: Before cryo-conservation at −80 °C the RNAlater was removed completely. Cartilage samples were cryo-pulverized (59012N, Biospec, Bartlesville, OK) and gently resuspended in TriFast™ (VWR, Germany) mixed with 1-bromo-3-chloropropane (Sigma Aldrich, MO). Centrifugation was performed after 10 min of incubation for 10 min at 10.000 × *g.* The top aqueous phase was further used for RNA isolation using the RNeasy Mini Kit (QIAGEN, Germany) according to the protocols provided. cDNA synthesis and qPCR were performed as described above.

### Statistical analysis

Statistical analysis was performed with the GraphPad Prism V.8 software. Quantitative data is shown as mean ± SEM for gene expression data and mean ± SD for all other data. Since the number of samples was small and Gaussian distribution was not assumed, we used the Kruskal-Wallis test with Dunn’s multiple comparison as non-parametric statistical test for group differences. Each SFCC from one donor was assumed to be an individual replicate, and thus paired analysis was not performed. Statistical numbers and adjusted p-values are listed in Supplementary Results. A p-value of < 0.05 was considered to be statistically significant. Image analysis was carried out blinded for treatment groups. Important numbers and adjusted p-values are stated either in the text or in the graphs.

### In silico model generation

To describe the temporal evolution of the spatial distribution of cellular and matrix-related processes in the *in vitro* model of OA, PDEs were used to interpret the experimental outcomes and to enable the identification of key steps within the progression of matrix degradation. The parameters of the *in silico* model were calibrated based on the histomorphometric *in vitro* data for distribution of collagen II (Col-2) and cell numbers. Since each section of the SFCCs which were analyzed had a thickness of 8 µm, we transformed the dimension from area (mm^2^) to volume (m^3^). The geometrical shape of the SFCCs was approximated by a cylindrical form (radius r = × = 5.0·10^−3^ m and height h = y = 4.5·10^−3^ m) (Figs. 1 and 5A). This cylinder-like construct was surrounded by culture medium, so that the added IL-1β was able to diffuse into the SFCC from all surfaces. The cylindrical form of the SFCC was radially symmetric around its center. Therefore, the computations were performed on a two-dimensional rectangle instead of the full three-dimensional domain (Figs. 1 and 5A). In this first modeling approach, we only focused on the effects of IL-1β, although the *in vitro* model was additionally stimulated with TNFα. The generated systems of time-dependent PDEs were solved using the state-of-the-art software package KARDOS [25, 26]. Given a user-specified accuracy tolerance, the code automatically computes numerical approximations by means of a fully adaptive grid in space and time based on local error estimates with optimal computational complexity. The graphical outputs were produced using MATLAB (Version 2017b, The MathWorks, Inc., Natick, MA).

## Results

### SFCCs show stable cartilage-like phenotype over 3 weeks

To evaluate the cellular and matrix composition of the SFCCs under non-inflammatory conditions, samples were taken weekly over a period of 3 weeks (d0, 7, 14, 21; n = 3; one sample per donor at each time point). Alcian blue staining for SFCC sections was performed at all time points and revealed the constant presence of glycosaminoglycans (GAGs) (Fig. 3A). Hematoxylin & eosin (H&E) staining showed no obvious morphological changes in cell and matrix composition between time points (Fig. 3A). Cell density decreased over the period of 3 weeks from a mean cell density of 1.378 ± 694 cells/mm^2^ at d0 to a mean cell density of 505 ± 268 cells/mm^2^ at d21 (Fig. 3B). In addition, histomorphometry revealed a higher cell count in the outer area (Ot.A.) than it did in the total area (Tt.A.) (Fig. 2 and Table 2), indicating a spatial arrangement of cells within the construct (Fig. 3B). Collagen I (Col-1) was more abundantly present within the constructs compared to Col-2 although no difference could be identified between time points (Fig. 3C). qPCR analyses revealed the mRNA expression of the cartilage specific markers *COL2A1* and *ACAN*, although *COL1A1* expression was higher when compared to that of *COL2A1* (Fig. 3D). *COL10A1* expression was low in comparison with all other genes. No statistical differences were found between time points. To conclude, SFCCs show a cartilage-like phenotype, although Col-1 does remain present.

**Table 4:**
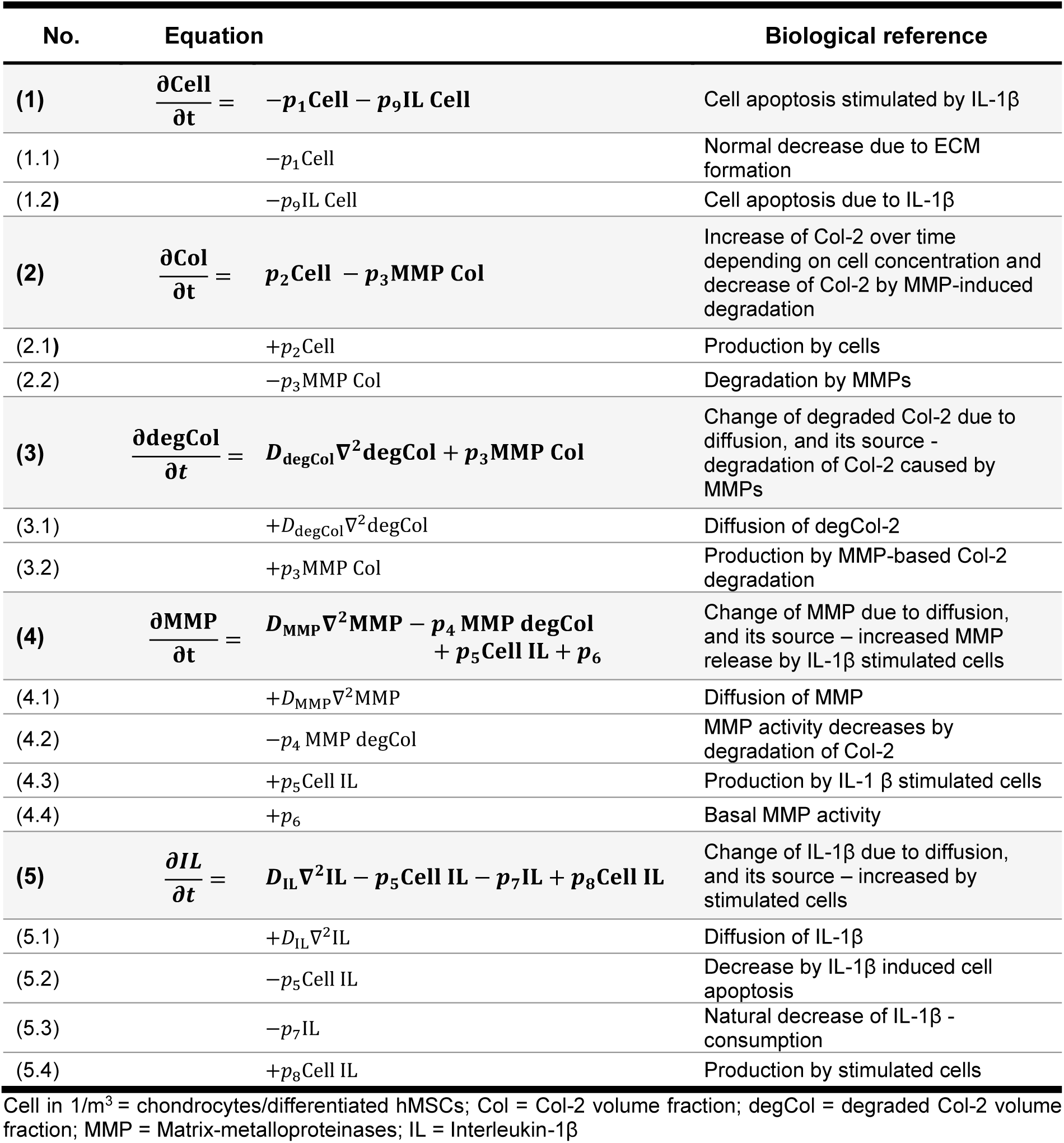
PDE model of the *in vitro* model resembling the onset of OA based on Kar et al. [16]

**Table 5:** Reduced PDE model of *in vitro* model resembling onset of OA

| No. | Equation | Biological reference |
| --- | --- | --- |
| **(1)** | $\frac{\partial \text{Cell}}{\partial t} = -p_1 \text{Cell} - p_9 \text{IL Cell}$ | Cell apoptosis stimulated by IL-1β |
| **(2)** | $\frac{\partial \text{Col}}{\partial t} = -p_3 \text{MMP Col}$ | Decrease of Col-2 by MMP-induced degradation |
| **(3)** | $\frac{\partial \text{MMP}}{\partial t} = D_{\text{MMP}} \nabla^2 \text{MMP} + p_5 \text{Cell IL}$ | Change of MMP due to diffusion and increased MMP release by IL-1β stimulated cells |
| **(4)** | $\frac{\partial \text{IL}}{\partial t} = D_{\text{IL}} \nabla^2 \text{IL}$ | Diffusion of IL-1β |

### Stimulated SFCCs show cellular and matrix-related changes comparable to those in the early phase of OA

To model the inflammatory environment of OA, SFCCs were treated with IL-1β and TNFα. both of which are considered to be the most important proinflammatory cytokines at the onset of OA. Cytokines were applied with concentrations of 50 ng/ml for IL-1β, and 100 ng/ml for TNFα representing a highly aggressive proinflammatory stimulation in order to achieve maximal effects on the SFCCs. Tissue softening was observed macroscopically by volume increase of SFCCs stimulated for 3 weeks (STIM) compared to untreated controls (CTL) which was partly reversed after 3 additional weeks without cytokine treatment (REG) (Fig. 4A). This observation was supported by histological H&E staining, microscopically indicating pronounced water retention, maceration of the superficial cell layer and morphological changes in cell phenotypes in the STIM group (Fig. 4B). Immunohistological analysis showed a constant Col-1 coverage with no statistically significant difference between groups (Fig. 4C). Col-2 coverage however decreased after stimulation with IL-1β and TNFα to a mean coverage of 14.4 ± 5.5% (STIM) and moreover significantly to 8.8 ± 5.9% (REG), when compared to the untreated controls (25.3 ± 4.7%; Fig. 4C). Histomorphometry revealed statistically significant changes between groups in cell count per area for the Tt.A., Ot.A. and outer core area (Ot.C.A.; Fig. 4D). Therefore, a lower cell density was observed in the STIM group compared to the CTL, while slight changes were also seen in the REG group compared to CTL. No significant differences were found in the In.C.A. (Fig. 4D). On the mRNA level, *COL1A1* was significantly downregulated upon cytokine stimulation in comparison with the control (Fig. 4E). The gene expression of *COL2A1* and *ACAN* was obviously lower in the STIM and REG group. Interestingly, *COL10A1* was upregulated in the REG group (Fig. 4E). As expected, gene expression levels of the inflammatory markers *IL1, IL6* and *IL8* were significantly upregulated compared to untreated controls (Fig. 4F). Gene expression of *TNF* was not found to be different between the experimental groups. However, *IL1, IL6* and *IL8* were numerically diminished in the REG compared to the STIM group although no statistical significance was detected (Fig. 4F). *MMP1* and *MMP3* were significantly upregulated in the cytokine-treated group compared to the CTL group and numerically downregulated in the REG compared to the STIM group (Fig. 4G). There were no significant differences in gene expression for *MMP13* within the experimental groups. In summary, stimulation with IL-1β and TNFα leads to OA-like changes which can be observed *in vivo* during the early phase of the disease. Additional cultivation for 3 weeks without cytokines after stimulation did partially reverse those changes.

### Refining an existing mathematical model results in a reduced PDE model of the in vitro observations to simulate the onset of OA

In order to focus on the main processes of cartilage degradation expected during OA onset, the *in silico* model was built on the assumption that under non-inflammatory conditions, the chondrocytes (termed here as cells) produce ECM (here mainly Col-2) while under inflammatory conditions with the addition of IL-1β, cells release MMPs, which then degrade the ECM, and potentially go into apoptosis (see Figure 1). Therefore, Table 4 shows the underlying equations derived from the pathway described above and our *in vitro* observations (adapted from [16]).

Considering the underlying *in vitro* experiments and the complexity of the derived *in silico* model, it was necessary to derive a reduced model preserving our *in vitro* observations and the underlying biology. The reduced model (Table 5) was obtained as described in the following. For equation (1), it was possible to distinguish between the decrease due to the matrix formation and due to IL-1β by comparing the reduction of the cell density under non-inflammatory and inflammatory conditions in the *in vitro* models. Thus, we could identify the parameters *p*_1_ (non-inflammatory conditions) and *p*_9_ (inflammatory conditions) separately. Following Kar et al., we expected a small increase of Col-2 under non-inflammatory conditions (small value for *p*_2_) which was negligible compared to the significant decrease induced by the MMPs [16]. Furthermore, we have omitted equation (3) for degCol-2. This component cannot be reliably measured and has no significant impact on other components. MMP measurements are only used for verification (Fig. 1), and therefore, parameter *p*_4_ and the basal MMP activity *p*_*6*_ – for which no estimates exist, have been omitted. Finally, in the IL-1β equation (5), the diffusion from outside is certainly the predominant source of IL-1β, which can also be verified e.g. by performing simulations of the model of Kar et al. with and without a correspondingly simplified equation for IL-1β.

In Table 6, we prescribed the parameter values, initial values and the boundary conditions to solve the PDE system. Concerning the parameter values, the cell apoptosis rate *p*_1_ was fitted by a least squares approach, utilizing measurements of the cell concentration (n=3, d= 0, 7, 14, 21). Since the MMP measurements have been neglected up to now, the diffusion coefficient is taken from the literature. The remaining parameters have been calibrated with our *in vitro* observations. For MMPs, we assumed that there are homogeneous Neumann boundary conditions for all surfaces. Because the evolution of the cell number and Col-2 was described by ODEs, i.e., where no diffusion is present, there exist no boundary conditions for these components. Due to the diffusion of IL-1β into the SFCC with a constant concentration of IL-1β in the surrounding media, inhomogeneous Dirichlet boundary conditions were assumed to be present on the outer surfaces. For the construct’s inner surface (center of the cartilage construct), homogeneous Neumann boundary conditions were applied ensuring the symmetry assumption on the SFCC.

**Table 6:**
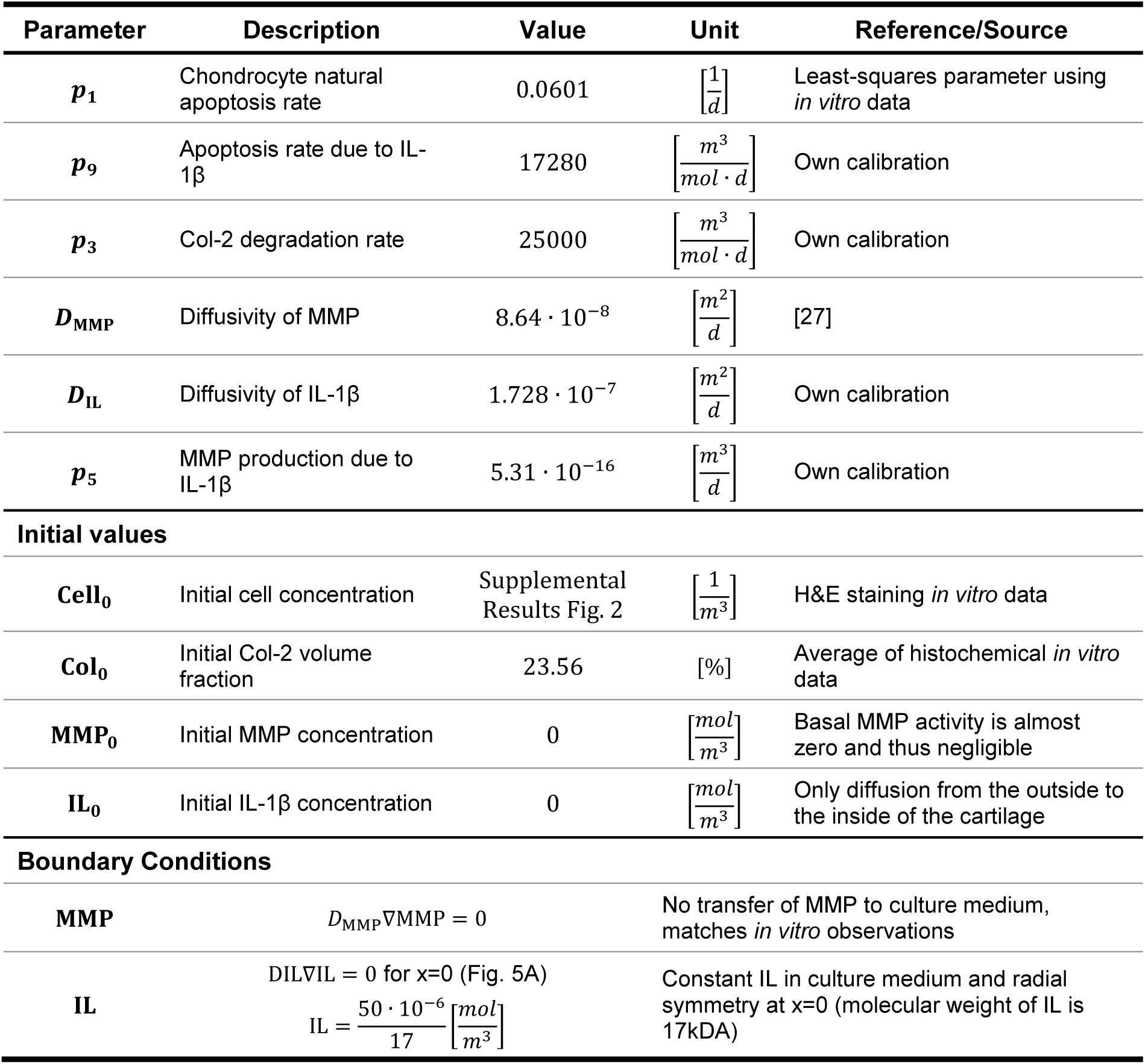
Parameter list, initial values and boundary conditions for reduced PDE model as in Table 5.

### Modeling the in vitro findings of cytokine-stimulated SFCCs in silico to resemble basic processes of OA cartilage degradation

Since the chondrocytes are inhomogeneously distributed over their space (Fig. 3), we determined an initial distribution based on the *in vitro* observations (Supplementary Fig. S1). Computational results of the *in silico* model are shown in Figure 5. The model described in Table 5 makes it possible to represent the *in vitro* experiments. Under IL-1β stimulation, the cell number decreased over time given the initial distribution (first row of Fig. 5B), which matches the *in vitro* measurements (Fig. 4D). Col-2 decreased spatiotemporally assuming a homogeneous initial distribution (second row in Fig. 5B). Since IL-1β enters from the outside and stimulated the MMP production, MMPs increased on the boundaries first and diffused further into cartilage (third row in Fig. 5B and Fig. 5C). Under non-inflammatory conditions, the IL-1β concentration is assumed to be zero. This also holds true for MMP (see Table 5, equations (3) and (4)). Thus, Col-2 will be constant over time and cells decrease only due to matrix formation leading to a representation of the healthy state observed *in vitro* as well.

To predict cellular and matrix-related changes over a longer time period (5 weeks) and a 10-fold lower concentration of IL-1β stimulation (5 ng/ml) – which is more comparable to the *in vivo* situation – we accordingly adapted the *in silico* model. We observed a full penetration of IL-1β and MMPs, a fast decline in the cell number, but a rather slower Col-2 degradation when compared to the 3 weeks simulation which had higher IL-1β concentrations (Supplementary video).

### MMP staining in the SFCCs verifies in silico simulations

In order to verify the mathematical model, MMP-1 and MMP-13 were stained via immunofluorescence for CTL, STIM and REG sections and analyzed for their spatial distribution and displayed as a sum of that. Therefore, MMP normalization to the cell number was slightly more visible in the STIM group (Fig. 5D). The relative MMP coverage was lower in the inner core area (In.C.A.) than it was in that of the Ot.C.A. Such a finding applies to a match between the CTL with the STIM groups and also among themselves (Figs. 5E, F). This also matches the results gained from the *in silico* model and nicely verifies our approach.

## Discussion

Since multiple approaches and a variety of OA model systems are necessary to fully understand the complexity of this disease, we report here on a successful combination of *in vitro* and *in silico* modelings to simulate the main features of OA, inflammation and cartilage destruction through upregulation of matrix degrading enzymes.

In most cases, it is only cartilage explants or monolayer cell cultures which are used as an *in vitro* model to study OA or to test potential therapeutic targets. However, there are some studies using tissue-engineered cartilage for their experiments such as those of Mohanraj et al. who report it to show the same features as cartilage explants in a model of posttraumatic OA [8]. [28]. In addition, there are some comparable approaches which focus on the production of SFCCs [9, 11, 29-31]. SFCCs revealed a cartilage-like phenotype detected by gene expression and histological analysis (Fig. 3); however, significant differences do remain when those are compared to native human cartilage (Supplementary Fig. S2). Thus, the gene expression of *COL1A1* and the presence of Col-1 on a protein level were still visible in all SFCCs examined. hMSCs have the potential to differentiate into different lineages including the chondrogenic, osteogenic and adipogenic lineage, although it has not been yet described that MSCs can fully form human hyaline articular cartilage *in vitro* or *in vivo*. Nevertheless, hMSCs are capable to form cartilage-like tissues which have many similarities to *in vivo* articular cartilage. Since the 3D structure of the SFCCs was gained due to biomechanical loading, the application of transforming growth factor beta (TGF-β) would not have been as beneficial as had been shown previously [32].

The *in vitro* simulation of the inflammatory environment in OA through a cytokine treatment with IL-1β at 50 ng/ml and TNFα at 100 ng/ml allowed us to see significant changes in the SFCCs towards an OA phenotype (Fig. 4). Although the *in vivo* concentrations for IL-1β and TNFα are known to be much lower (e.g. as demonstrated in synovial fluid of patients suffering from knee OA [33]), cytokine concentrations of 1-100 ng/ml are usually applied when mimicking the proinflammatory environment in early OA *in vitro.* The latter situation is necessary in order to shorten the periods of time for treatment by adapting the *in vitro* experiments, because the chronic degenerative joint disease OA in humans usually evolves over decades. IL-1β and TNFα do not only contribute to the upregulation of proteases but also inhibit the synthesis of ECM molecules, mainly Col-2 and aggrecan [34, 35]. MMPs constitute one predominant group of enzymatic proteins which plays an important role in the pathogenesis of OA [36, 37].

Mathematical models are very flexible, enabling adjustments and testing of various hypotheses in parallel. However, biological experiments are most often quite complex, expensive, resource-demanding and risky, leading to a gain of less experimental data than model parameters can [38]. Thus, non-identifiability problems can occur, since model parameters cannot be estimated properly. Experimental design approaches aim to resolve these problems by identifying the data gap and proposing the required additional experimental data. The flexibility of mathematical models including experimental design can be used to structure future *in vitro* or *in vivo* models efficiently. However, the first considered mathematical model, which takes most of the prominent mechanisms of OA pathogenesis into account, has unidentifiable parameters (Table 4). Its applicability is nevertheless limited, since neither enough experimental human data nor reliable literature-based parameters are available for model calibration and validation. This means that more quantitative measurements of the components at different time and spatial points are needed. The reduced model, however, does allow us to reproduce the experimental data with significantly fewer parameters (Table 5). Validation concerning MMP measurements which were not used to calibrate the model indicates that the reduced *in silico* model is suited to investigate the influence of different targets for the early phase of OA (Fig. 5). In addition, the Supplementary video in this article shows the *in silico* model’s feasibility as a prediction tool for further *in vitro* experiments. The observed fast decline in the cell number accompanied by a slower Col-2 degradation is in accordance with a previously published study indicating a high chondrocyte death before matrix changes occur [39]. In comparison to the model described by Kar et al. [16], our model considers changes of the cell concentration in space as well as over time. A more detailed analysis of the model proposed by Kar et al. revealed IL-1α as the driving force in the model, while MMP, Col-2 and degCol-2 only have a small influence on the remaining components. Thus, the simplified model in our study here displays the same features but at the same time involves a smaller number of parameters and uncertainties. Other mathematical simulation approaches for OA and cartilage focus on e.g. poroelastic models or coupling cellular phenotype and mechanics [19, 40, 41].

In summary, we here describe a human 3D *in vitro* model of OA based on SFCC which simulates the main features of cartilage destruction, inflammation and upregulation of matrix degrading enzymes. Advantages of such a model include the 3D environment, the possibility for mid-throughput, a sufficient sample volume for several analyses and a wide availability of tissue-engineered cartilage with more scaffold-free approaches emerging. With the combination of *in vitro* and *in silico* modeling and close cooperation between biologists and mathematicians, we aim at paving the way towards new, innovative approaches which allow the immediate adaptation and modifications of different models. Within this unique collaboration, we strived to develop a mathematical model to refine and optimize the *in vitro* model and vice versa, especially regarding the development of a whole joint *in vitro/in silico* model.

### Limitations

The proposed *in vitro* model displays only one specific part of the disease and includes only one tissue type, although OA has been described as a whole organ disease including several tissues such as the synovial membrane and the subchondral bone. Thus, we are currently working on a more complex whole joint model for both *in vitro* and *in silico* aspects. Secondly, biomechanical loading has not yet been addressed within the first *in vitro* and *in silico* models presented here, although one’s awareness of its tremendous impact has already been published [42]. Thirdly, we shortened the experimental time window by using high concentrations of the proinflammatory cytokines IL-1β and TNFα in our experiments. That bridges the long-lasting cumulative effect of these cytokines over years and decades in the course of OA pathogenesis. Finally, an extension of the *in silico* model should include the treatment with TNFα due to its playing an important role in OA by describing the disease pathogenesis more accurately. For a profound model validation and parameter fitting, more quantitative *in vitro* data will be necessary.

## Supporting information

Supplementary Results

## Acknowledgments

The authors would like to thank Manuela Jakstadt for her excellent technical assistance. We also thank Kar et al. for making their mathematical model available for our study. The scientific discussions with David Smith (The University of Western Australia) and Atte Eskelinen (University of Eastern Finland) were very helpful and greatly appreciated. AL, FB, AD, MP and TG are members of Berlin-Brandenburg research platform BB3R and Charité 3^R^.

## Contributions

AL, SR, FB and RE designed the study. MCW, AD, MP, TG, FB and AL collected, analyzed and interpreted data on the *in vitro* model. IP generated SFCCs. LF, SG, JL, SR and RE developed mathematical model and performed simulations. AL, RE, MCW, AD and LF discussed and optimized mathematical model. MCW, LF, SG, RE and AL prepared the main manuscript text. All authors contributed to writing or reviewing the manuscript and final approval.

