## Supplementary Results for "*In vitro* and *in silico* modeling of cellular and matrix-related changes during the early phase of osteoarthritis"

**Table S1:** Results from the Kruskal-Wallis test for Figure 2

| Figure 2 | Specification | Kruskal-Wallis test | |
|---|---|---|---|
| | | H | *p*-value |
| B | Cells – Tt.A. 95% | 6.39 | *0.79* |
| | Cells – Ot.A. 5% | 0.15 | *0.99* |
| C | Col-1 | 5.77 | *0.12* |
| | Col-2 | 1.97 | *0.63* |
| D | *COL1A1* | 2.93 | *0.62* |
| | *COL2A1* | 7.45 | *0.09* |
| | *ACAN* | 2.23 | *0.74* |
| | *COL10A1* | 4.57 | *0.37* |

**Table S2:** Results from the Kruskal-Wallis test for Figure 3

| Figure 3 | Specification | Kruskal-Wallis test | |
|---|---|---|---|
| | | H | *p*-value |
| C | Col-1 | 2.68 | *0.27* |
| | Col-2 | 8.59 | *0.006* |
| D | Tt.A. 95% | 10.71 | *0.001* |
| | Ot.A. 5% | 9.36 | *0.004* |
| | Ot.C.A. 50% | 10.33 | *0.002* |
| | In.C.A. 50% | 5.07 | *0.075* |
| E | *COL1A1* | 6.49 | *0.012* |
| | *COL2A1* | 4.62 | *0.1* |
| | *ACAN* | 5.60 | *0.05* |
| | *COL10A1* | 5.96 | *0.03* |
| F | *IL1* | 7.2 | *0.004* |
| | *IL6* | 7.2 | *0.004* |
| | *IL8* | 6.49 | *0.01* |
| | *TNF* | 0.36 | *0.88* |
| G | *MMP1* | 6.49 | *0.01* |
| | *MMP3* | 7.2 | *0.004* |
| | *MMP13* | 1.16 | *0.63* |

**Table S3:** Results from the Dunn's multiple comparisons test

| Specification | Dunn's multiple comparisons test | |
|---|---|---|
| | **Comparison** | **Adjusted *p*-value** |
| **Fig. 3C Col-2** | CTL vs. STIM | *0.20* |
| | STIM vs. REG | *0.68* |
| | CTL vs. REG | *0.01* |
| **Fig. 3D Tt.A. 95%** | CTL vs. STIM | *0.005* |
| | STIM vs. REG | *0.99* |
| | CTL vs. REG | *0.05* |
| **Fig. 3D Ot.A. 5%** | CTL vs. STIM | *0.007* |
| | STIM vs. REG | *0.48* |
| | CTL vs. REG | *0.30* |
| **Fig. 3D Ot.C.A. 50%** | CTL vs. STIM | *0.008* |
| | STIM vs. REG | *0.99* |
| | CTL vs. REG | *0.03* |
| **Fig. 3E *COL1A1*** | CTL vs. STIM | *0.03* |
| | STIM vs. REG | *0.41* |
| | CTL vs. REG | *0.89* |
| **Fig. 3F *IL1*** | CTL vs. STIM | *0.02* |
| | STIM vs. REG | *0.54* |
| | CTL vs. REG | *0.54* |
| **Fig. 3F *IL6*** | CTL vs. STIM | *0.02* |
| | STIM vs. REG | *0.54* |
| | CTL vs. REG | *0.54* |
| **Fig. 3F *IL8*** | CTL vs. STIM | *0.03* |
| | STIM vs. REG | *0.41* |
| | CTL vs. REG | *0.89* |
| **Fig. 3G *MMP1*** | CTL vs. STIM | *0.03* |
| | STIM vs. REG | *0.41* |
| | CTL vs. REG | *0.89* |
| **Fig. 3G *MMP3*** | CTL vs. STIM | *0.02* |
| | STIM vs. REG | *0.54* |
| | CTL vs. REG | *0.54* |

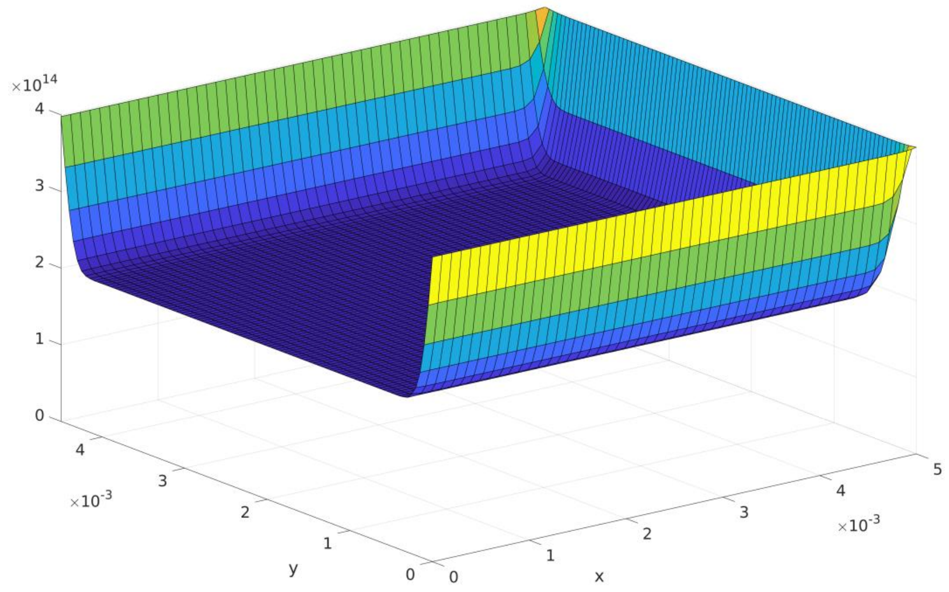


**Figure S1: Initial cell distribution** [$1/m^3$], see Fig. 5A for an explanation of the dimensions and axes.

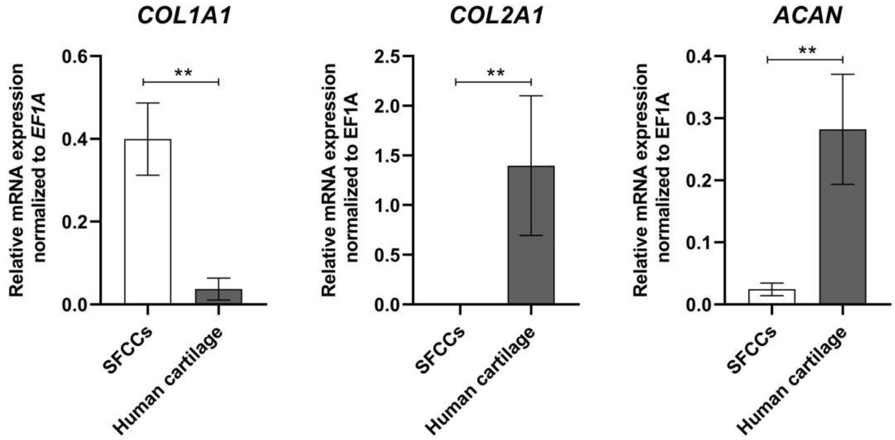


**Figure S2: RNA expression of COL1A1, Col2A1 and ACAN in SFCCs compared to native human cartilage.** Relative mRNA expression (2^-ΔCt) normalized to the housekeeper *EF1A* graph bar with mean ± SEM. For SFCCs all data from d0-21 were grouped together (Fig. 2D). Cartilage was collected from three different donors from almost unaffected sites (femoral condyles). Statistical differences between groups was tested with the Mann-Whitney test. **p < 0.01

**Supplementary Video: Using the *in silico* model as prediction tool.** The video displays the changes over time when applying 50 ng/ml IL-1β for 3 weeks (left side) as done in the *in vitro* experiments compared to the changes that are expected (based on the developed PDE model) when supplementing a 10fold lower concentration of IL-1β for 5 weeks. Cell [1/m$^3$], Col-2 [%], IL-1$\beta$ [mol/m$^3$] and MMP [mol/m$^3$].
